# Identification and kinetics characterization of a wax ester hydrolase from a feather-degrading actinomycete

**DOI:** 10.1101/178673

**Authors:** Matthew Barcus, Dario Mizrachi, Xin Gen Lei

## Abstract

*Streptomyces fradiae* var. k11 is a Gram-positive soil microorganism capable of degrading chicken feathers. Apart from being mostly protein, chicken feathers have a considerable level of lipids, with wax esters being the largest lipid class. The waxes may pose a challenge while rendering the feathers into coproducts, such as feather meal, and so the identification of a wax-ester hydrolase is warranted. A draft genome sequence of *S. fradiae* var. k11 was used to identify 14 gene sequences of potential lipid-degrading enzymes. The genes were expressed in *E. coli* BL21(DE3) cells on a pET vector and screened for activity. Four of the 14 enzymes had detectable activity, with two of the enzymes, SFK3309 and SFK3087, active against p-nitrophenyl palmitate, a representative water-insoluble substrate. A modified enzymatic assay was designed to measure activity against three model wax substrates: jojoba oil, beeswax, and cetyl-palmitate. SFK3309 was characterized to hydrolyze all three wax substrates. Kinetic experiments for SFK3309 were performed with cetyl-palmitate at 37°C, pH 8.0. The *K*_*m*_ was determined to be 850 µM and the *K*_*cat*_ was 11.63 s^-1^. Through the characterization of SFK3309 as a wax-ester hydrolase, biotechnological implications of wax ester hydrolases in the rendering of many industrial wastes can be substantiated for further studies.

## Introduction

Chicken feathers are well known for being predominately protein (90%), mostly comprised of recalcitrant ß-keratin (Gupta et al. 2012). However fat content has been known to vary inversely with protein content in feather meal, anywhere from 2-12% (Dale 1992). Some groups have estimated chicken feather fat to be near 11% (Kondamudi et al. 2009). Chickens secrete lipids from the uropygial gland onto their plumage for protection from the elements in an act known as preening. Wax (di)esters account for 49.3% of total preen oil composition and 38.2% of total feather lipids (Wertz et al. 1986).

Feathers represent approximately 3-6% of a chicken’s whole body weight (Leeson and Walsh 2004) and with the annual slaughter of 8.9 billion chickens in the USA (USDA (U.S. Department of Agriculture) 2017), effective rendering of chicken feathers is necessary to combat pollution. The protein from chicken feathers has considerable economic value, but the significant presence of fat in chicken feathers may pose a challenge for adequate degradation of the ß-keratin, and thus lower the economic potential.

Over half a century ago, *Streptomyces fradiae* was identified for the biodegradation of wool and feather keratin at enriched waste sites (Noval and Nickerson 1959) and our lab has recently been investigating enzymes from *S. fradiae* var. k11 for industrial relevance. Already the research community has identified many keratinases and reductases in *S. fradiae* (Gupta and Ramnani 2006), but we wanted to determine if lipolytic enzymes capable of hydrolyzing the waxy feather lipids were also present. Multiple strains of *Streptomyces* have been previously characterized to utilize beeswax found on ancient artwork as a sole carbon source (Sakr et al. 2013). Given that *S. fradiae* var. k11 is of the same genus, it may be reasonable to suspect *S. fradiae* var. k11 also produces enzymes capable of catalyzing the hydrolysis of wax esters found on chicken feathers.

A wax ester is a fatty acid esterified to a fatty alcohol and is critical for multiple physiological events. Wax esters are used by some plants and marine organisms as an energy storage (Leray 2006). Often on the epicuticular or epidermal layers of plants and animals, wax esters provide defense against pathogens and desiccation (Leray 2006). Wax esters are also a buoyancy regulator in fish (Phleger 1998). The unique properties of wax esters have significant industrial potential for use in cosmetics (Schlossman and Shao 2014), pharmaceuticals (Coffey 2012; El Mogy 2004), art, food products (Hepburn et al. 2000), fuel (Al-Widyan and Al-Muhtaseb 2010), and as a lubricants (Bhatia et al. 1990; Kalscheuer and Steinbuchel 2003). There have been multiple groups that have focused on the biosynthesis of wax esters (Heilmann et al. 2012; Rodriguez et al. 2014; Santala et al. 2014; Trani et al. 1991; Tsujita et al. 1999), but studies focused on the hydrolysis of wax esters have been few. The literature contains a few examples of wax ester hydrolases (EC 3.1.1.50) from plant (Huang et al. 1978; Kalinowska and Wojciechowski 1985; Moreau and Huang 1981), fungal (Brahimihorn et al. 1989), marine (Benson et al. 1975; Kayama et al. 1979; Mankura et al. 1984; Patton et al. 1975), and bacterial (Kalscheuer and Steinbuchel 2003) origins have previously been reported, however kinetic information is scant (Huang et al. 1978; Kalinowska and Wojciechowski 1985).

The identification and characterization of wax ester hydrolases has utility in the development of biotechnology to process agricultural byproducts with waxy surface lipids, such as wool (Brahimihorn et al. 1991) and feathers. Moreover, the similar physical characteristics of waxes and plastics (solid, insoluble) has facilitated in the discovery of a wax worm digesting plastic bags (Bombelli et al. 2017) or cutinase enzymes (EC: 3.1.1.74) hydrolyzing polyethylene terephthalate (PET) (Guebitz and Cavaco-Paulo 2008). Our report identified, and characterized, one wax ester hydrolase from a feather-degrading actinomycete with wax ester hydrolase activity.

## Materials and Methods

### Strains and chemicals

Chemicals were purchased from Sigma Aldrich (St. Louis, Missouri, USA) unless otherwise noted. Bacteria strain *Streptomyces fradia* var. k11 was gifted to the lab from collaborators (Meng et al. 2007).

### Cloning of wax esterase genes

*S. fradiae* var. k11 was cultured in brain heart infusion broth and genomic DNA was isolated as described previously by Nicodinovic, et al (Nikodinovic et al. 2003). Previous work in the lab has yielded a draft genome for *S. fradiae* var. k11. The genome was annotated on the Rapid Annotation using Subsystem Technology (RAST) server using the RASTClassic annotation scheme to identify potential lipolytic enzymes (Aziz et al. 2008). Additional enzymes were identified with patterns PDOC00110, PDOC00842, and PDOC00903 from the PROSITE database (Hulo et al. 2006). Some of the partial sequences were completed using sequence information from Ju et al. (Ju et al. 2015) and the previously reported *lips221* sequence (Zhang et al. 2008) was downloaded from GenBank (ID: EF429087.1) (Benson et al. 2013). Genomic DNA served as template for a polymerase chain reaction (PCR) to amplify out the genes. Pred-TAT (Bagos et al. 2010), an online signal peptide recognition tool, was used to predict the presence of any signal sequence. Oligonucleotides were designed to pull out the different genes without any native signal peptide or stop codon, while incorporating NcoI and XhoI restriction sites at the 5’ and 3’ ends, respectively.

The PCR reaction was performed using Platinum pfx polymerase purchased from Thermo Fisher Scientific (Waltham, Massachusetts, USA). The placement of restriction sites allowed for simplified cloning of PCR products into the pET22b(+) and pET28a vectors via double digestion. Ligated constructs were cloned into *E. coli* BL21(DE3) cells and selected over LB amp^+^ plates. Plasmid DNA was isolated using a QIAGEN (Venlo, Netherlands) Qiaprep Spin miniprep kit and the construct was verified at the Cornell Biotechnology Resource Center DNA Sequencing facility using T7 oligonucleotides. Sequencing results were analyzed against a vector map using ApE - A Plasmid Editor (Wayne Davis, Salt Lake City, Utah, USA; version 2.0.49.10) (Davis 2008).

### Gene expression and protein purification

Starter cultures in Luria broth (LB) were grown overnight and used to inoculate expression cultures. Expression cultures used either LB, terrific broth (TB), or autoinduction media (Teknova, Hollister, California, USA) and were grown at 37^o^C to OD_600_ of 0.6-0.8, then placed into a 16°C, shaking incubator for 22 hours. Cultures in LB or TB were induced with 0.1 mM Isopropyl β-D-1-thiogalactopyranoside (IPTG).

Protein was harvested by first collecting the cells via centrifugation (5000 × *g* for 5 minutes). Cells were resuspended in lysis buffer (20 mM HEPES, 500 mM NaCl, 10 mM imidazole, pH 7.6). Cells were then sonicated for 8 minutes using 4 second pulses at 45% amplitude on a Sonics (Newtown, Connecticut, USA) Vibracell sonicator (model: VC 130). The disrupted cells were then centrifuged at 15,000 × *g* for 20 minutes and the total soluble protein was mixed with Ni-NTA agarose (McLab, South San Francisco, California, USA) for 1 hour on a rolling shaker at 4°C. The slurry was then loaded into a gravity flow column and allowed for the agarose to settle. Upon the agarose settling, the column was uncapped and the flow-through was collected. The Ni-NTA agarose was then washed and collected three times with wash buffer (20 mM HEPES, 500 mM NaCl, and 20 mM imidazole, pH 7.6). Protein bound to the Ni-NTA agarose was then eluted with elution buffer (20 mM HEPES, 500 mM NaCl, and 250 mM imidazole, pH 7.6).

The eluted protein was then concentrated in an Amicon ultra-15 centrifuge tube with a molecular weight cut-off of 10 kDa. Concentrated protein then further purified using an ÄKTA Explorer FPLC system (GE Healthcare, USA). Size exclusion chromatography (SEC) was performed on all NiNTA purified proteins. The column employed for purification was Superdex 200 10/300 GL. Standards used to calibrate the SEC column were a lyophilized mix of thyroglobulin, bovine γ-globulin, chicken ovalbumin, equine myoglobin, and vitamin B12 (Molecular weight range: 1,3500-670,000 Da, pI range: 4.5-6.9) (Bio-Rad, Hercules, California, USA). Proteins were stored at 4°C in SEC buffer (20 mM HEPES, 500 mM NaCl, 5 mM CaCl_2_, pH 7.6) until further analyzed for activity and purity. Protein purity was determined by sodium dodecyl sulfate - polyacrylamide gel electrophoresis (SDS-PAGE) using Coomassie blue staining (Simpson 2007).

### Enzyme activity assays

Protein concentration was determined as previously described (Walker 2009), with the exception of the incubation taking place at 37°C instead of 60°C. Lipase assays were conducted using p-nitrophenyl palmitate as substrate. Assay conditions have been previously described (Zhang et al. 2008). Briefly, p-nitrophenyl palmitate (pNPP) was dissolved in 2-propanol to make a 10 mM solution. A reaction buffer was prepared (50 mM sodium phosphate, pH 8.0, 0.1% gum arabic, 0.2% sodium deoxycholate) and mixed with the pNPP solution in a 9 to 1 ratio. Enzyme sample (10 µL) was loaded into a 96-well microtiter plate and 240 µL of the substrate solution was added immediately before taking a kinetic read for 2 minutes with 15 second intervals using a wavelength of 410 nm on a SpectraMax M2e spectrophotometer (Molecular Devices, Sunnyvale, California, USA). One enzyme unit was defined as the hydrolysis of 1.0 µmole of p-nitrophenyl palmitate per minute at pH 8.0 at room temperature. Other p-nitrophenyl ester substrates (butyrate, decanoate, and stearate) were used interchangeable with pNPP for the assay.

Wax ester hydrolase activity was determined using three different wax substrates: beeswax, cetyl palmitate, and jojoba oil. Wax substrate was solubilized in 2-propanol, using a heat block when necessary and mixed with the same reaction buffer from the pNPP assay and at the same ratio. The substrate solution was then added to a microcentrifuge tube that contained the enzyme sample and was incubated in a 37°C, shaking incubator for up to 1 hour. Released fatty acid products from wax ester hydrolysis were then detected using reagents from the NEFA-HR(2) kit from Wako Chemicals (Richmond, Virginia, USA). Briefly, in a 96-well microtiter plate an 8 µL sample from the enzyme reaction was mixed with 144 µL color reagent A, which contained coenzyme A (CoA), acyl-CoA synthetase, and adenosine triphosphate and resulted in the formation of acyl-CoA. After a 5-minute incubation at 37°C, the microtiter plate was read at A_550_. Color reagent B (acyl-CoA oxidase, peroxidase, 3-methyl-ethyl-N-(ß-hydroxyethyl)-aniline(MEHA), and 4-aminoantipyrine) was then added (48 µL), which oxidized the acyl-CoA into hydrogen peroxide, thus allowing MEHA to undergoing oxidative condensation with 4-aminoantipyrine and create a purple colored product. After a 5-minute incubation at 37°C, a second A_550_ reading was made. A standard curve was created using varying concentrations of oleate and an enzyme unit was defined as the release of 1 millimole-equivalent free fatty acid per minute per milligram of enzyme.

Thin layer chromatography (TLC) was performed as described previously (Yamada et al. 2016). Briefly, an enzyme reaction with a wax substrate was set up as described earlier. The reactions were then incubated in a 37°C shaking incubator for 1 hour. Methanol (100 µL) was added to the sample and vortexed. Chloroform (200 µL) was then added and the sample was again vortexed. A sample collected from the chloroform layer was used to spot a HPTLC silica gel 60 TLC plate and double-developed with eluent hexane: diisopropyl ether: acetic acid (50:50:1). The plate was dried and then sprayed with coloring agent (27.4 mM 12-molybdo(VI) phosphoric acid n-hydrate and 50 mM sodium perchlorate dissolved in ethanol). The plate was then heated in an oven for 2 minutes until the bands were revealed. A separate reaction was set up with a heat-deactivated enzyme and was performed concurrently with the reaction containing the active enzyme. The enzyme was heated to 65°C for 10 minutes to fully deactivate it.

Data analysis was carried out using the Pandas module in python (McKinney 2011). Plots were generated using Matplotlib (Hunter 2007).

## Results

### Identification of lipolytic enzymes from *Streptomycete* genome

From a draft genome sequence of feather-degrading actinomycete *S. fradiae* var. k11, fourteen genes were identified as potential lipases from annotations and ProSite patterns. Four genes, out of the pool of 14, had detectable enzymatic activity against p-nitrophenyl acetate or pNPP (Table 1). Lipase activity was observed only in two of the four identified enzymes (SFK3087 and SFK3309), as demonstrated by their ability to hydrolyze pNPP. Both SFK3309 and SFK3087 were tested against the wax ester substrate jojoba oil, but only SFK3309 retained activity and further characterization of SFK3309 was warranted. **Sequence characterization of SFK3309**

**Table 1.**
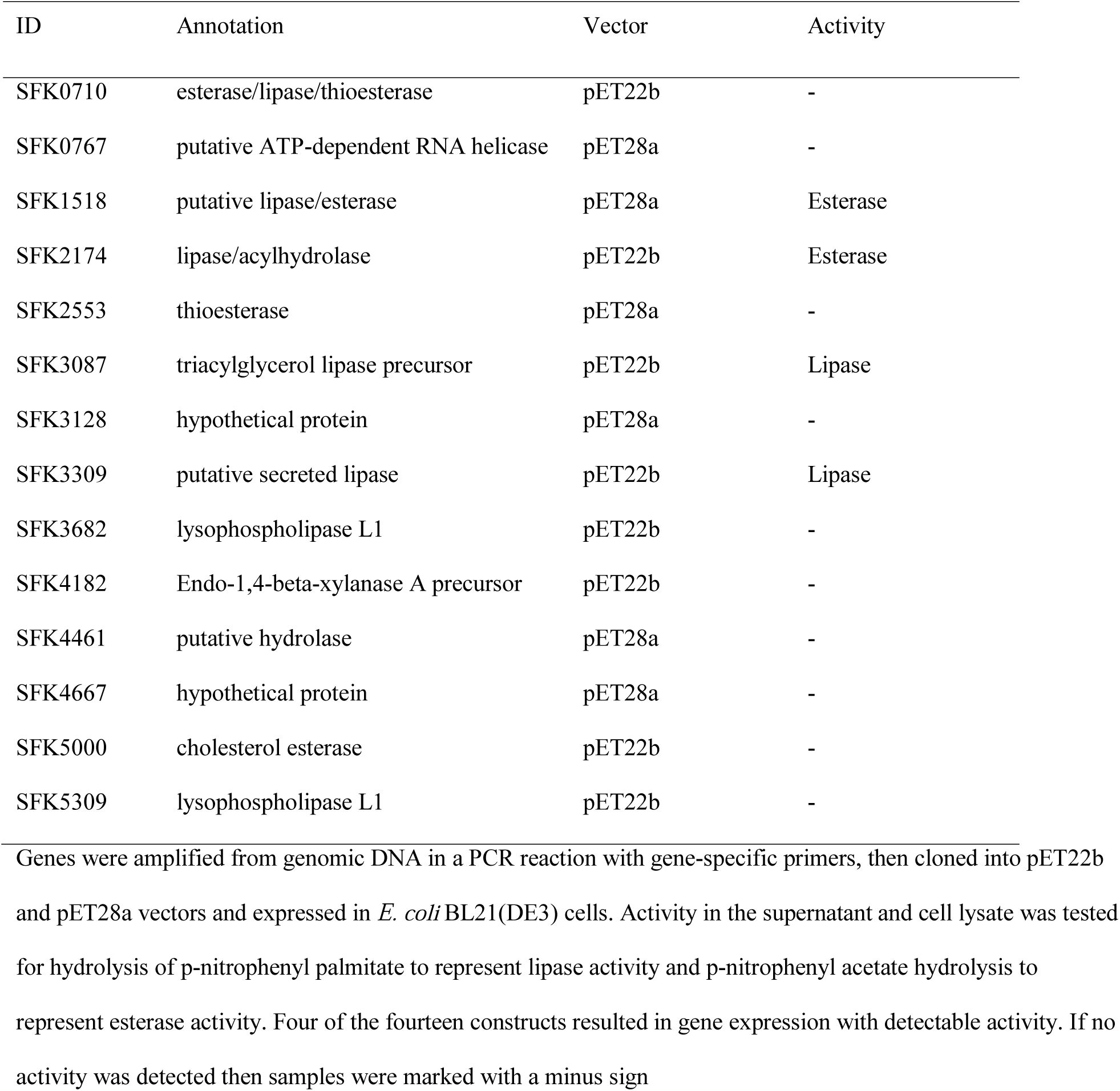
Lipolytic activity screen for genes from *Streptomyces fradiae* var. k11

The cloned construct of SFK3309 in the pET22b(+) was verified via Sanger sequencing. Gene sequences of SFK3309 and lips221 which served as the gene template for cloning were aligned (Fig. 1). The translated sequence codes for four alternative residues from lips221: P86L, L88P, N223S, and L237P.

**Fig. 1.**
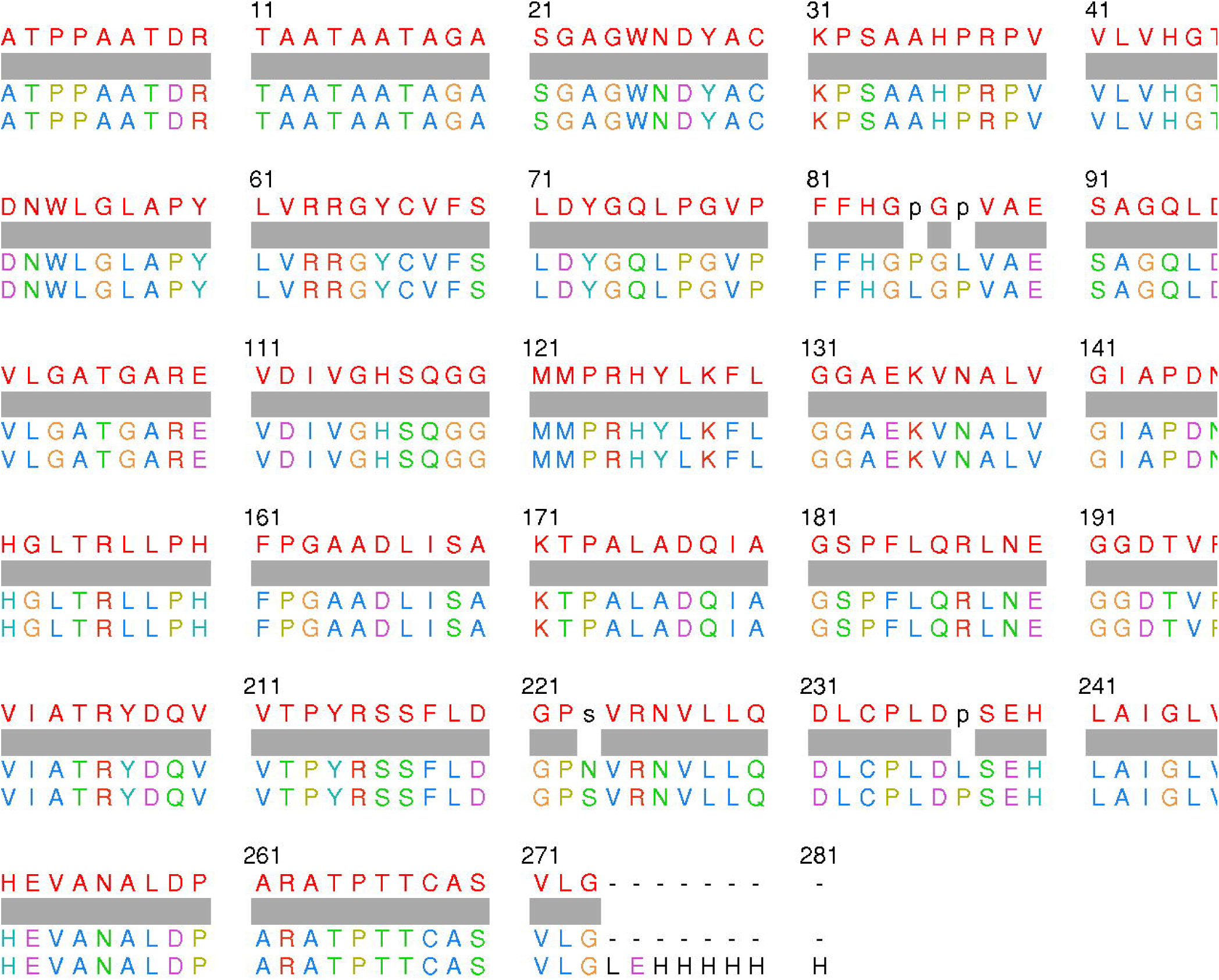
SFK3309 alignment with lips221. Using MAFFT (Katoh and Standley 2013) the top performing lipase from this study (SFK3309) was aligned to lips221 (GenBank ID: EF429087.1), a highly similar sequence initially reported by Zhang, et al (Zhang et al. 2008). Four missense mutations were present between the two sequences: P86L, L88P, N223S, and L237P. Alignment was visualized in Chimera (Pettersen et al. 2004)

### Expression optimization and purification

Initially, culturing pET22b SFK3309 in *E. coli* BL21(DE3) cells appeared to have some toxicity as indicated by a reduced OD_600_ at harvest time compared to that of the time of IPTG induction. Cultures expressing SFK3309 were tested using different growth media for optimization: Luria broth (LB), terrific broth (TB), and Studier’s ZYM-5052 autoinduction media (AT). Not only did the cell density of cultures grown in AT increase compared to LB and TB (OD_600_ ∼6.0), the enzyme activity in the lysate was ∼3X higher, as seen in Figure 2.

**Fig. 2.**
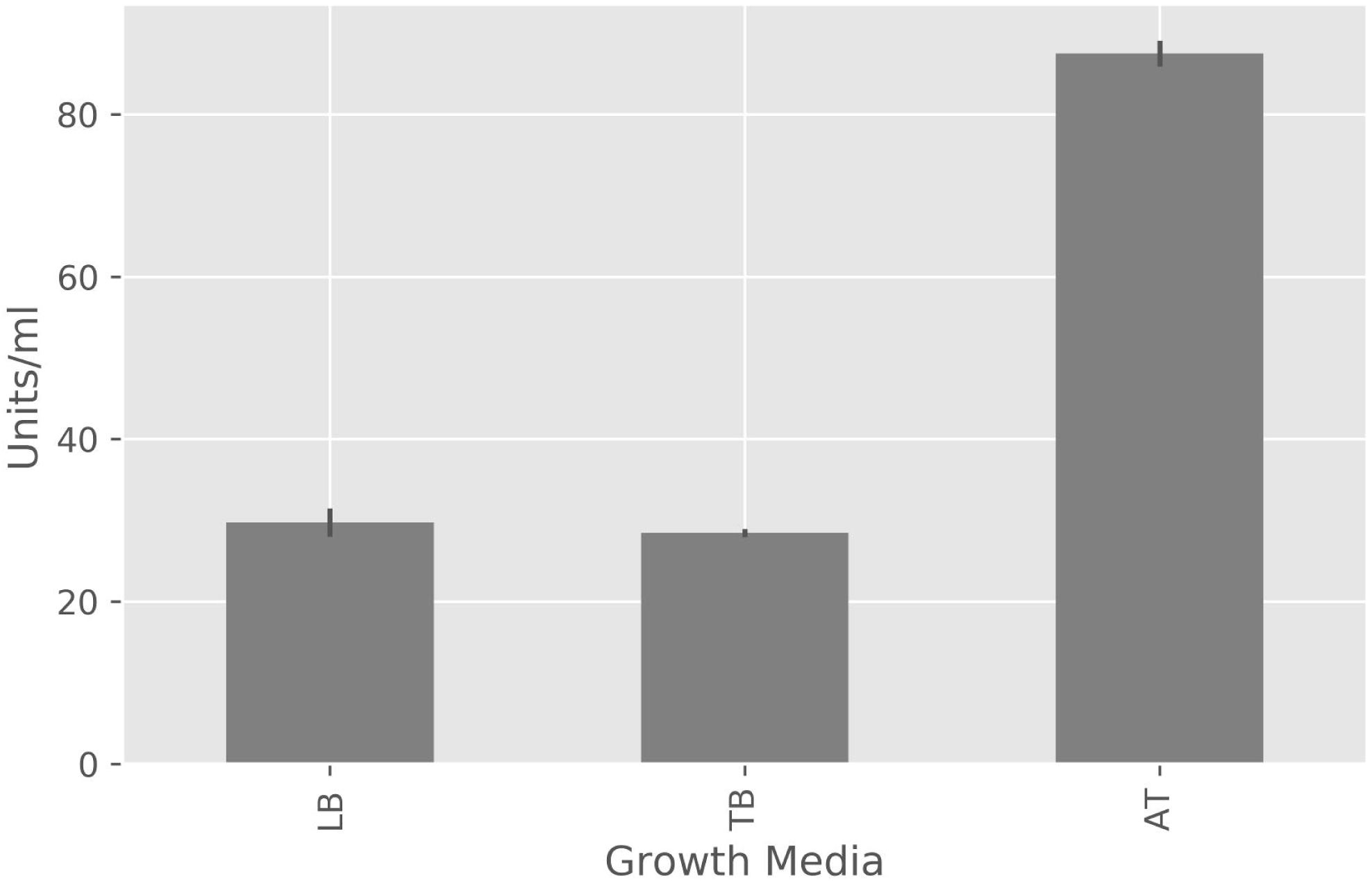
Optimal growth media for SFK3309. Luria broth (LB), Terrific broth (TB), and Studier's ZYM-5052 autoinduction media (AT) were tested for impact on protein expression. Cultures were grown as described in *Materials in Methods*. Assays were performed in duplicate and the standard deviation is represented by the error bars. Both LB and TB had comparable activity in the lysate ∼22 hours post IPTG induction while AT had 2.9X the activity in the lysate at the same time period

The C-terminal His tag on the pET vector was used for the first purification step of SFK3309. After eluting SFK3309 from the Ni-NTA agarose column, the protein was run on an SDS-PAGE gel and stained using Coomassie blue (Fig. 3). A subsequent gel filtration step was used and separated the multimeric protein forms.

**Fig. 3.**
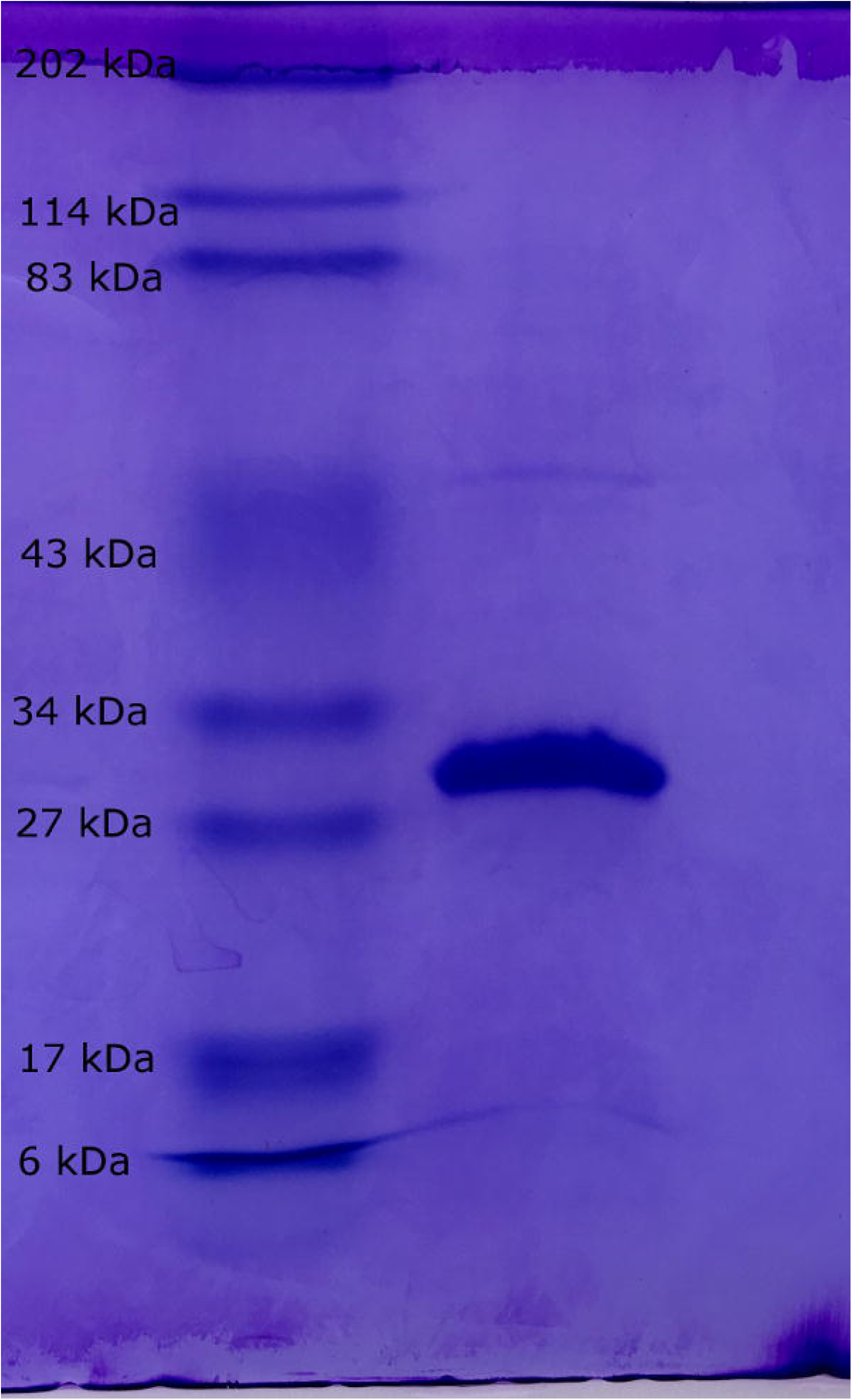
Coomassie blue stained SDS-PAGE gel of his-tag purified SFK3309. The left lane is the Bio-Rad broad range protein marker and its bands are labelled with their respective molecular weights. The right lane is the purified SFK3309 (29.6 kDa) eluted from the Ni-NTA agarose (see Materials and Methods)

### Enzyme activity analysis

Substrate chain-length preference of SFK3309 was determined by the release of nitrophenol from synthetic substrates with varying length carbon-chains esterified to nitrophenyl. Substrates included: para nitrophenyl-butyrate (C4), para nitrophenyl-decanoate (C10), para nitrophenyl-palmitate (C16), and para nitrophenyl-stearate (C18). Preference of SFK3309 for the shorter carbon chain of para nitrophenyl-butyrate was evident (Fig. 4).

**Fig. 4.**
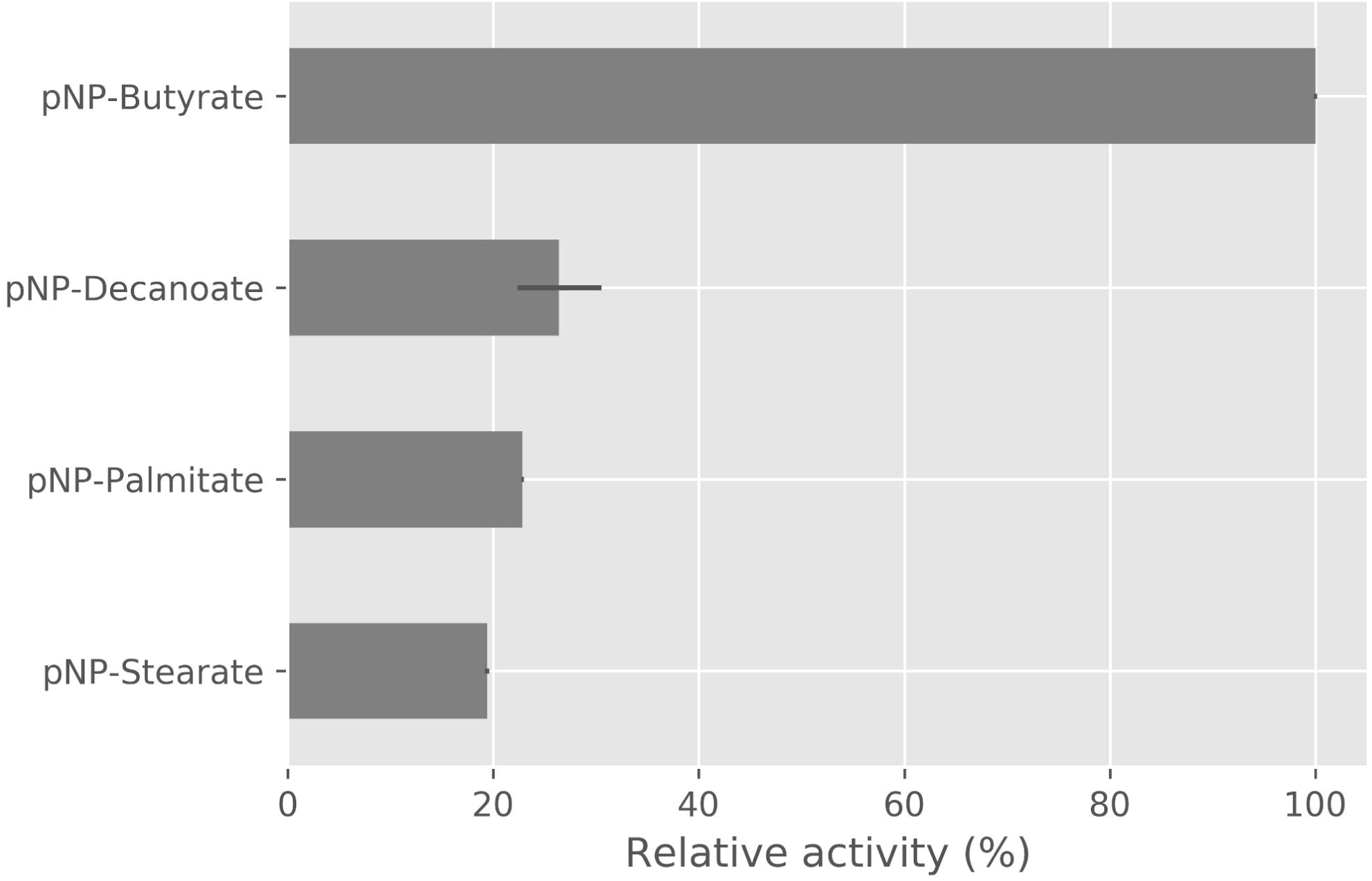
Substrate chain-length preference of SFK3309. Purified SFK3309 was evaluated for enzyme hydrolysis activity against the chromophore p-nitrophenol (pNP) esterified to different carbon-chain length carboxylates: butyrate (C4), decanoate (C10), palmitate (C16), and stearate (C18). Reaction conditions were carried out at room temperature buffered with 0.05 M sodium phosphate, pH 8.0 and performed in duplicate. The average activity of SFK3309 against pNP-Butyrate was considered absolute (100%) and the standard deviation was represented by the error bars. The data suggests SFK3309 has preference for the shorter-chained substrate, yet retains broad specificity, allowing the enzyme to retain activity against all the substrates tested

Kinetic studies of SFK3309 with cetyl-palmitate as substrate were performed. A Lineweaver-Burk plot was used for the determination of the *K*_*m*_ and *V*_*max*_ values (Fig. 5). The *K*_*m*_ was worked out to be 8.5x10^−3^ M and the *K*_*cat*_ was 11.55 s^-1^, yielding a *K*_*cat*_*/K*_*m*_ value of 1.35 • mMs^-1^ (Table 2). Visual hydrolysis of cetyl-palmitate, alongside two more substrates, jojoba oil and beeswax, was carried out using thin layer chromatography (Fig. 6).

**Fig. 5.**
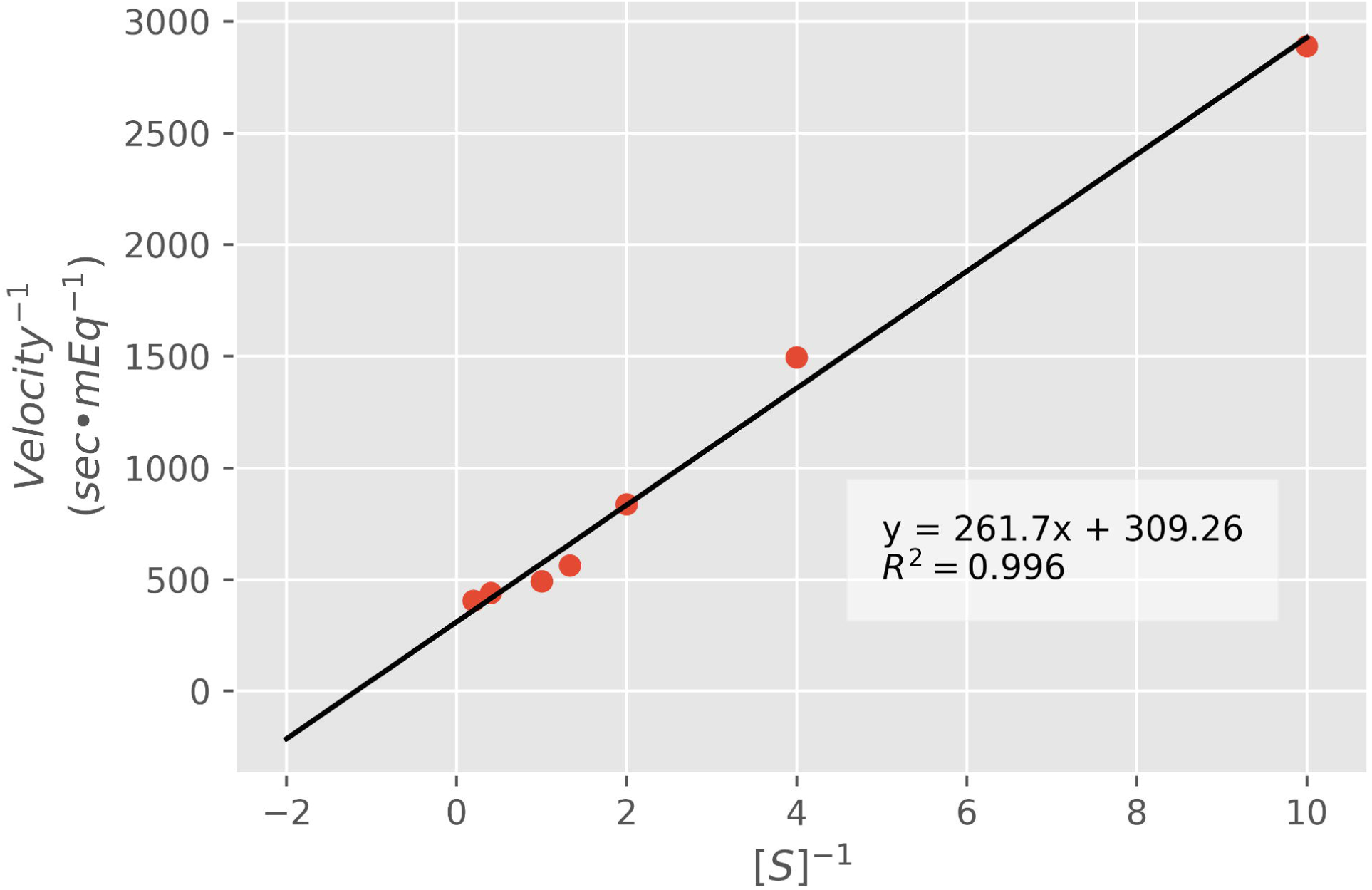
Lineweaver-Burk plot for kinetics calculations of cetyl-palmitate hydrolysis by SFK3309. Cetly-palmitate concentrations ranged from 0.1 to 5 mM. Reactions were carried out at 37°C for 5 minutes. The concentration of SFK3309 was 0.206 mg/mL. All reactions were performed in triplicate and the average values were plotted. The *K*_*cat*_ was determined to be 11.63 s^-1^ and the *k*_*cat*_*/k*_*m*_ was 13.74/mM•s^-1^. The plot was made in python using Matplotlib (Hunter 2007)

**Table 2.** – Comparison of reported wax-ester hydrolysis kinetics values.

| $K_m$ (M) | $K_{cat}$ (1/s) | $K_{cat}/K_m$ (1/mMs<sup>-1</sup>) | Organism | Reference |
| --- | --- | --- | --- | --- |
| $2.8 \times 10^{-5a}$ | -- | -- | *Sinapis alba* L. | (Kalinowska and Wojciechowski 1985) |
| $9.3 \times 10^{-5b}$ | -- | -- | *Simmondsia chinensis* | (Huang et al. 1978) |
| $8.5 \times 10^{-4c}$ | 11.63 | 13.7 | *S. fradiae* var. k11 | Current study |

<sup>a</sup> Radiolabeled cetyl-palmitate was used as substrate in this study.
<sup>b</sup> N-methylindoxyl myristate was used as substrate.
<sup>c</sup> Unlabeled cetyl-palmitate was used as substrate.

**Fig. 6.**
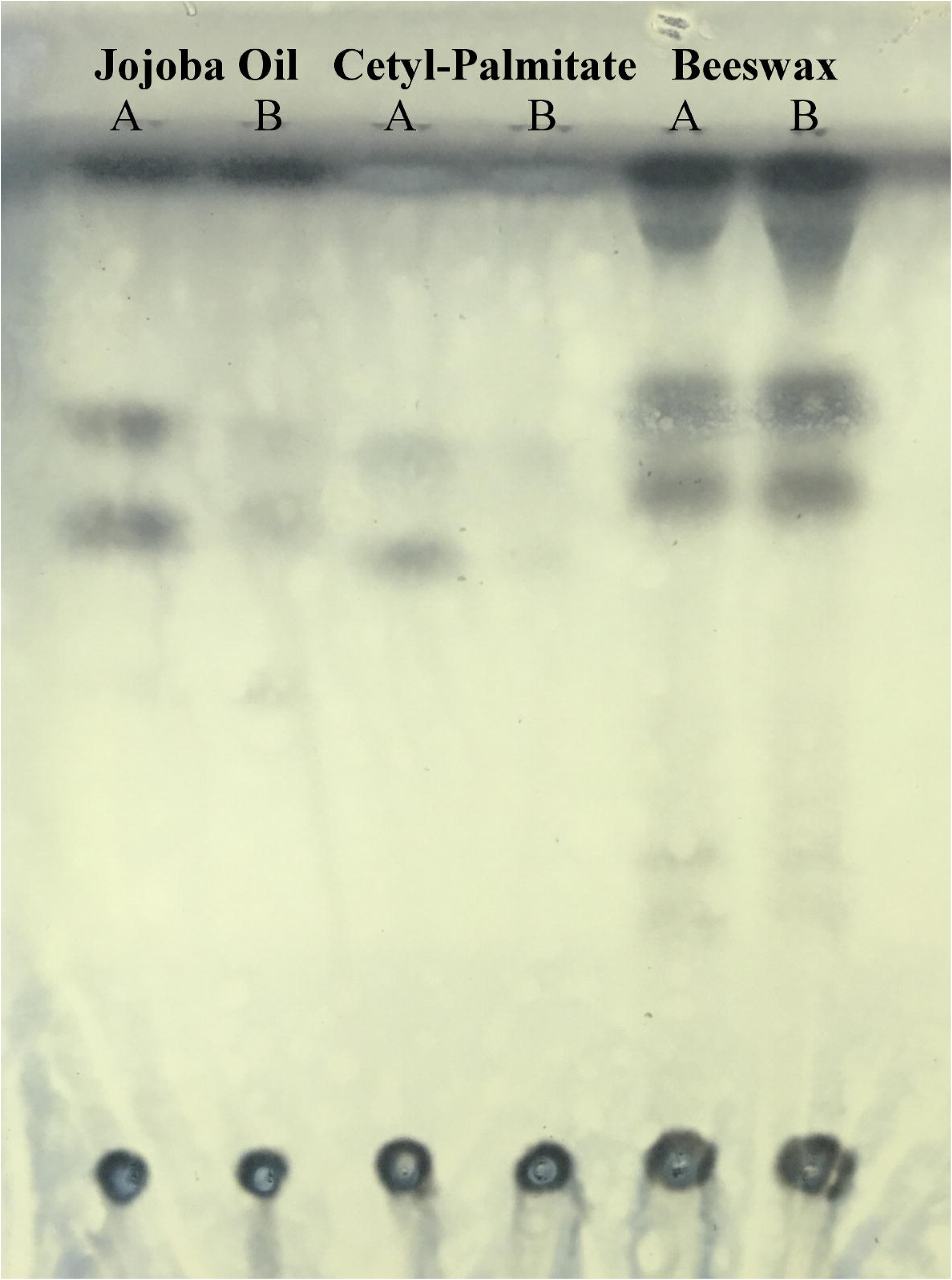
Thin layer chromatography of SFK3309 wax hydrolysis products. Wax-ester hydrolase assays were set up with substrates jojoba oil, cetyl-palmitate, and beeswax and treated with active SFK3309 (A) or SFK3309 that had been heat-deactivated (B). The samples with active SFK3309 appeared to create more products than the heat-deactivated SFK3309 against jojoba oil and cetyl-palmitate as represented in the two thicker lower bands, though the differences in beeswax were negligible

## Discussion

An initial set of 14 gene candidates were identified in feather-degrading bacterial strain *S. fradiae* var. k11 to encode for lipolytic activity. From this set, four enzymes expressed successfully in *E. coli* BL21 (DE3) and had detectable activity towards pNPA or pNPP (Table 1). The simplicity of the pNPP and pNPA assays allowed for a rapid identification of active enzymes and permitted the differentiation of enzymes capable of hydrolyzing long-chain, water-insoluble substrates (pNPP) or only active towards short-chained water-soluble substrates (pNPA). As the physical properties of wax suggest, it was necessary to identify the enzymes capable of acting on the water-insoluble substrate. Two enzymes were capable of hydrolyzing pNPP: SFK3087 and SFK3309.

The Lipase Engineering Database (LED) (Fischer and Pleiss 2003) classified SFK3087 as part of the alpha-beta hydrolase superfamily 25, while the ESTHER database (Lenfant et al. 2013) classified the enzyme as a member of the lipase class 1 and polyesterase-lipase-cutinase family. SFK3309 was classified by LED to the alpha-beta hydrolase 16 superfamily and the ESTHER database suggested the enzyme belongs to the lipase class 2 family. Given SFK3087 was classified as a member of the polyesterase-lipase-cutinase family, it seemed likely the enzyme would perform well against longer-chained substrates, such as plant cutin. However, only SFK3309 was active against wax substrates, and not SFK3087. Therefore, SFK3309 was selected as the central enzyme of the study.

DNA sequence analysis of SFK3309 cloned in the pET22b vector indicated four missense mutations (P85L, L87P, N223S, and L237P) from the lips221 sequence, as shown in the MAFFT (Katoh and Standley 2013) alignment (Fig. 1). Secondary structure prediction of lips221 by JPRED4 (Drozdetskiy et al. 2015) suggested all SFK3309 mutations fell in loop regions and are less probable to severely disrupt overall protein structure.

A pelB signal sequence on the pET22b vector directed SFK3309 to the periplasm to provide an ideal environment for secretory protein expression (Boock et al. 2015). Expression of SFK3309 was carried out in *E. coli* BL21 (DE3) cells. Lipolytic activity in the supernatant as well as a decrease in final OD_600_ suggested some level of toxicity from expressing SFK3309 in the cells, a familiar issue in heterologous lipase expression (Drouault et al. 2000). Three different growth media were evaluated for improved SFK3309 expression: LB, TB, and Studier’s ZYM-5052 media. LB is standard media found in labs, while TB provides added glycerol and uses a phosphate buffer (Boock et al. 2015), an understandable advantage considering free fatty acid products from lipolysis can affect pH. Studier’s ZYM-5052 media is also buffered, but was able to further improve protein titers through using a blend of carbon sources and bypassing the need for IPTG induction for stable expression (Studier 2005). Expression experiments demonstrated LB and TB activity in the cell lysate was comparable, while Studier’s ZYM-5052 medium improved expression levels about 3-fold (Fig. 2).

By using the C-terminal 6X-histidine tag in the pET22b vector SFK3309 was purified from the cell lysate and able to reach approximately 90% purity from one round of Ni-NTA purification as demonstrated from SDS-Page (Fig. 3). Yields of purified protein for 250 mL cultures were about 2 mg. A subsequent size-exclusion chromatography step was used to further separated SFK3309 from different oligomeric states. While Zhang et al. struggled with *E. coli* BL21 (DE3) as a heterologous host for extracellular expression of lipS221 (Zhang et al. 2008), our experience suggests reasonable protein expression levels can be achieved in *E. coli* BL21 (DE3) for SFK3309 by coupling periplasmic expression with autoinduction media. The reported *K*_*m*_ of 0.139 mM of lips221 expressed in *Pichia pastoris* (Zhang et al. 2008) was comparable to the *K*_*m*_ of 0.078 mM that our group produced with SFK3309 against pNPP. The *K*_*cat*_ of SFK3309 was determined to be 1668 s^-1^, which was within reason of values reported by similar enzymes (Dheeman et al. 2011).

SFK3309 was evaluated for wax ester hydrolase activity using three different substrates: jojoba oil, beeswax, and cetyl palmitate. Jojoba oil is derived from the seeds of *Simmondsia chinensis,* a plant native to the southwestern United States. The major lipid form in jojoba oil are wax esters, with the dominant carbon chain length being C_40_ and C_42_ (Bussonbreysse et al. 1994). Jojoba oil was an ideal substrate to work with initially due to the low melting point (10°C) allowing for easy handling and the negligent levels of free fatty acids made the detection of reaction products unambiguous. Beeswax is produced by honey bees and is 35% composed of wax monoesters with mostly C_40_ – C_48_ chain lengths (Tulloch 1971) and myricyl-palmitate as the main component (Riemenschneider and Bolt 2005). The higher melting temperature (63°C) of beeswax required heating in order to dissolve into 2-propanol, and the raw substrate further tested enzyme utility. Cetyl-palmitate was the third wax substrate used for analysis of wax-ester hydrolase activity. Cetyl-palmitate, like beeswax is a solid at room temperature (melting point = 54°C) and thus required similar handling. However, given cetyl-palmitate is easily obtained as a pure substrate, wax-ester hydrolase kinetic experiments were possible.

Previous groups have reported wax ester hydrolase assays that rely on radioactively labelled substrates (Bogevik et al. 2008; Brahimihorn et al. 1989; Tsujita et al. 1999), which require resources not accessible to many labs. Others have also reported colorimetrically detecting free fatty acid products by first converting the fatty acids to copper salts and measured with sodium dithiocarbamate (Heinen and De Vries 1966; Huang et al. 1978) but these assays are time consuming and often rely on using strong organic solvents such as chloroform, thus limiting the use of plastic ware. The wax ester hydrolase assay used for this study first incubates an enzyme with a standard wax substrate, then utilizes the Wako NEFA HR(2) commercial kit to detect released free fatty acids. The entire assay can be performed in a basic lab equipped with a spectrophotometer using standard microtiter plates and microcentrifuge tubes and does not require an extraction step. The linear detection range of free fatty acids using the Wako NEFA HR(2) kit is 0.01-4.0 mEq/L. Since the standard curve is constructed using oleic acid, the millimole-equivalent unit is used to denote any free fatty acid released. Jojoba oil was used initially as a substrate to confirm the reliability of the assay.

SFK3309 exhibited enzyme activity against all three of the wax ester substrates using the described colorimetric assay. TLC was used to confirm the observed enzyme activity. From Figure 6, the active enzymes produced thicker lower bands, corresponding to the hydrolysis of the substrate, though beeswax was an anomaly, which may be due to a high level of background. Enzyme kinetics were determined using cetyl-palmitate as substrate. Two groups have previously reported on the kinetics of wax-ester hydrolases and their results are available for comparison in Table 2. In Huang et al.’s study the synthetic substrate N-methylindoxyl myristate was used for kinetic experiments of an enzyme from jojoba cotyledons (Huang et al. 1978). Their work found the enzyme’s *K*_*m*_ value to be 9.3 × 10^−5^ M, within close proximity to the work of Kalinowska and Wojciechowski, who used cetyl [1-^14^C]palmitate as substrate and reported a *K*_*m*_ of 2.8 × 10^−5^ M for an enzyme from the roots of white mustard (Kalinowska and Wojciechowski 1985). The constructed Lineweaver-Burk plot from our study suggests a good linear fit for our data (Fig. 5). The *K*_*m*_ of SFK3309 was determined to be 8.5 × 10^−4^, which is close to the other groups’ values. We also found the *K*_*cat*_ of SFK3309 to be 11.63 s^-1^

In conclusion, we identified two lipases from the feather-degrading bacteria *S. fradiae* var. k11 and characterize one of the enzymes as a wax-ester hydrolase. The described wax-ester hydrolase assay from our study allowed for the evaluation of enzyme activity against various wax substrates and permitted a kinetics experiment against cetyl-palmitate. Given the presence of waxy lipids on the surface of chicken feathers, the existence of wax-ester hydrolase activity from SFK3309 in *S. fradiae* var. k11 suggests a potential application in industrial feather hydrolysis.

Current literature on the enzymatic hydrolysis of wax esters is limited, with kinetic studies even more scarce. The adaptation of the wax ester hydrolase assay as reported here allows for a more accessible approach to evaluate wax ester hydrolase activity and yield results comparable with traditional techniques. Waxes are an important lipid class and have a presence across multiple industries. A more comprehensive investigation of enzymes capable of hydrolyzing wax esters may contribute to the development of existing and novel industrial processes.

## Acknowledgements

Dr. Shinya Fushinobu provided suggestions for the substrate specificity assay and TLC, as well as lab space. Dr. Jeremy Baskin and Dr. Robert Moreau for their input in design of the wax ester hydrolase assay. Vectors for heterologous expression in E. coli were provided by Dr. Matthew DeLisa.

## Compliance with Ethical Standards

### Funding

This work was supported by the National Science Foundation [grant number 1613943]; the Japan Society for the Promotion of Science [grant number SP160002]; and the Northeast Sustainable Agriculture Research and Education [grant number GNE13-052].

### Ethical Approval

This article does not contain any studies with human participants or animals performed by any of the authors.

## References

Al-Widyan MI, Al-Muhtaseb MA (2010) Experimental investigation of jojoba as a renewable energy source. Energ Convers Manage 51(8):1702–1707 doi:10.1016/j.enconman.2009.11.043

Aziz RK, Bartels D, Best AA, DeJongh M, Disz T, Edwards RA, Formsma K, Gerdes S, Glass EM, Kubal M, Meyer F, Olsen GJ, Olson R, Osterman AL, Overbeek RA, McNeil LK, Paarmann D, Paczian T, Parrello B, Pusch GD, Reich C, Stevens R, Vassieva O, Vonstein V, Wilke A, Zagnitko O (2008) The RAST server: Rapid annotations using subsystems technology. Bmc Genomics 9 doi:10.1186/1471-2164-9-75

Bagos PG, Nikolaou EP, Liakopoulos TD, Tsirigos KD (2010) Combined prediction of Tat and Sec signal peptides with hidden Markov models. Bioinformatics 26(22):2811–2817 doi:10.1093/bioinformatics/btq530

Benson AA, Patton JS, Field CE (1975) Wax Digestion in a Crown-of-Thorns Starfish. Comp Biochem Phys B 52(2):339–340 doi:10.1016/0305-0491(75)90075-9

Benson DA, Cavanaugh M, Clark K, Karsch-Mizrachi I, Lipman DJ, Ostell J, Sayers EW (2013) GenBank. Nucleic Acids Res 41(D1):D36–D42

Bhatia VK, Chaudhry A, Sivasankaran GA, Bisht RPS, Kashyap M (1990) Modification of Jojoba Oil for Lubricant Formulations. J Am Oil Chem Soc 67(1):1–7 doi:Doi 10.1007/Bf02631379

Bogevik AS, Tocher DR, Waagbo R, Olsen RE (2008) Triacylglycerol-, wax ester- and sterol ester-hydrolases in midgut of Atlantic salmon (Salmo salar). Aquacult Nutr 14(1):93–98 doi:10.1111/j.1365-2095.2007.00510.x

Bombelli P, Howe CJ, Bertocchini F (2017) Polyethylene bio-degradation by caterpillars of the wax moth Galleria mellonella. Curr Biol 27(8):R292–R293 doi:10.1016/j.cub.2017.02.060

Boock JT, Waraho-Zhmayev D, Mizrachi D, DeLisa MP (2015) Beyond the cytoplasm of Escherichia coli: localizing recombinant proteins where you want them. Insoluble Proteins: Methods and Protocols:79–97

Brahimihorn MC, Guglielmino ML, Sparrow LG (1989) Wax Esterase-Activity in a Commercially Available Source of Lipase from Candida-Cylindracea. J Biotechnol 12(3-4):299–306 doi:10.1016/0168-1656(89)90049-7

Brahimihorn MC, Mickelson CA, Guglielmino ML, Gaal AM, Sparrow LG (1991) Identification of Lipolytic-Activity in a Multitrophic Population Grown in Wool Scour Effluent. J Ind Microbiol 8(1):53–58 doi: 10.1007/Bf01575591

Bussonbreysse J, Farines M, Soulier J (1994) Jojoba Wax - Its Esters and Some of Its Minor Components. J Am Oil Chem Soc 71(9):999–1002 doi: 10.1007/Bf02542268

Coffey MJ (2012) Ophthalmic compositions with wax esters. Google Patents

Dale N (1992) True metabolizable energy of feather meal. The Journal of Applied Poultry Research 1(3):331–334

Davis M (2008) Ape: A plasmid editor.

Dheeman DS, Henehan GTM, Frias JM (2011) Purification and properties of Amycolatopsis mediterranei DSM 43304 lipase and its potential in flavour ester synthesis. Bioresource Technol 102(3):3373–3379 doi:10.1016/j.biortech.2010.11.074

Drouault S, Corthier G, Ehrlich SD, Renault P (2000) Expression of the Staphylococcus hyicus lipase in Lactococcus lactis. Appl Environ Microb 66(2):588–598 doi: 10.1128/Aem.66.2.588-598.2000

Drozdetskiy A, Cole C, Procter J, Barton GJ (2015) JPred4: a protein secondary structure prediction server. Nucleic Acids Res 43(W1):W389–W394 doi:10.1093/nar/gkv332

El Mogy NS (2004) Medical effect of Jojoba oil in the treatment of anal diseases. Google Patents

Fischer M, Pleiss J (2003) The Lipase Engineering Database: a navigation and analysis tool for protein families. Nucleic Acids Res 31(1):319–321 doi:10.1093/nar/gkg015

Guebitz GM, Cavaco-Paulo A (2008) Enzymes go big: surface hydrolysis and functionalisation of synthetic polymers. Trends Biotechnol 26(1):32–38 doi:10.1016/j.tibtech.2007.10.003

Gupta A, Kamarudin NB, Kee CYG, Yunus RBM (2012) Extraction of keratin protein from chicken feather. J Chem and Cheml Eng 6(8):732

Gupta R, Ramnani P (2006) Microbial keratinases and their prospective applications: an overview. Appl Microbiol Biot 70(1):21–33 doi:10.1007/s00253-005-0239-8

Heilmann M, Iven T, Ahmann K, Hornung E, Stymne S, Feussner I (2012) Production of wax esters in plant seed oils by oleosomal cotargeting of biosynthetic enzymes. J Lipid Res 53(10):2153–2161 doi:10.1194/jlr.M029512

Heinen W, De Vries H (1966) A combined micro-and semi-micro colorimetric determination of long-chain fatty acids from plant cutin. Archiv für Mikrobiologie 54(4):339–349

Hepburn P, Quinlan PT, Smith KW, Watts JV, van der Wielen RPJ (2000) Wax ester compositions. Google Patents

Huang AHC, Moreau RA, Liu KDF (1978) Development and Properties of a Wax Ester Hydrolase in Cotyledons of Jojoba Seedlings. Plant Physiol 61(3):339–341 doi:10.1104/pp.61.3.339

Hulo N, Bairoch A, Bulliard V, Cerutti L, De Castro E, Langendijk-Genevaux PS, Pagni M, Sigrist CJA (2006) The PROSITE database. Nucleic Acids Res 34:D227–D230 doi:10.1093/nar/gkj063

Hunter JD (2007) Matplotlib: A 2D graphics environment. Comput Sci Eng 9(3):90–95 doi:10.1109/Mcse.2007.55

Ju KS, Gao JT, Doroghazi JR, Wang KKA, Thibodeaux CJ, Li S, Metzger E, Fudala J, Su J, Zhang JK, Lee J, Cioni JP, Evans BS, Hirota R, Labeda DP, van der Donk WA, Metcalf WW (2015) Discovery of phosphonic acid natural products by mining the genomes of 10,000 actinomycetes. P Natl Acad Sci USA 112(39):12175–12180 doi:10.1073/pnas.1500873112

Kalinowska M, Wojciechowski ZA (1985) Characterization of Wax-Ester Hydrolase from Roots of White Mustard (Sinapis-Alba L) Seedlings. Acta Biochim Pol 32(3):259–269

Kalscheuer R, Steinbuchel A (2003) A novel bifunctional wax ester synthase/acyl-CoA: diacylglycerol acyltransferase mediates wax ester and triacylglycerol biosynthesis in Acinetobacter calcoaceticus ADP1. J Biol Chem 278(10):8075–8082 doi:10.1047/jbc.M210533200

Katoh K, Standley DM (2013) MAFFT multiple sequence alignment software version 7: improvements in performance and usability. Mol Biol Evol 30(4):772–780

Kayama M, Mankura M, Ikeda Y (1979) Hydrolysis and Synthesis of Wax Esters by Different Systems of Carp Hepatopancreas Preparation. J Biochem-Tokyo 85(1):1–6

Kondamudi N, Strull J, Misra M, Mohapatra SK (2009) A Green Process for Producing Biodiesel from Feather Meal. J Agr Food Chem 57(14):6163–6166 doi:10.1021/jf900140e

Leeson S, Walsh T (2004) Feathering in commercial poultry - I. Feather growth and composition. World Poultry Sci J 60(1):42–51 doi:10.1079/Wps20044

Lenfant N, Hotelier T, Velluet E, Bourne Y, Marchot P, Chatonnet A (2013) ESTHER, the database of the alpha/beta-hydrolase fold 41(D1):D423–D429 doi:10.1093/nar/gks1154

Leray C (2006) Waxes. Kirk-Othmer Encyclopedia of Chemical Technology

Mankura M, Kayama M, Saito S (1984) Wax Ester Hydrolysis by Lipolytic Enzymes in Pyloric Ceca of Various Fishes. B Jpn Soc Sci Fish 50(12):2127–2131

McKinney W (2011) pandas: a foundational Python library for data analysis and statistics. Python for High Performance and Scientific Computing:1–9

Meng K, Li J, Cao YA, Shi PJ, Wu B, Han XY, Bai YG, Wu NF, Yao B (2007) Gene cloning and heterologous expression of a serine protease from Streptomyces fradiae var.k11. Can J Microbiol 53(2):186–195 doi:10.1139/W06-122

Moreau RA, Huang AH (1981) [93] Enzymes of wax ester catabolism in jojoba. Method Enzymol 71:804–813

Nikodinovic J, Barrow KD, Chuck J-A (2003) High yield preparation of genomic DNA from Streptomyces. Biotechniques 35(5):932–936

Noval JJ, Nickerson WJ (1959) Decomposition of Native Keratin by Streptomyces-Fradiae. J Bacteriol 77(3):251– 263

Patton JS, Nevenzel JC, Benson AA (1975) Specificity of Digestive Lipases in Hydrolysis of Wax Esters and Triglycerides Studied in Anchovy and Other Selected Fish. Lipids 10(10):575–583 doi:10.1007/Bf02532720

Pettersen EF, Goddard TD, Huang CC, Couch GS, Greenblatt DM, Meng EC, Ferrin TE (2004) UCSF chimera - A visualization system for exploratory research and analysis. J Comput Chem 25(13):1605–1612 doi:10.1002/jcc.20084

Phleger CF (1998) Buoyancy in marine fishes: Direct and indirect role of lipids. Am Zool 38(2):321–330

Riemenschneider W, Bolt H (2005) Esters, Organic Ulmann's Encyclopedia of Industrial Chemistry. p 8676–8694

Rodriguez GM, Tashiro Y, Atsumi S (2014) Expanding ester biosynthesis in Escherichia coli. Nat Chem Biol 10(4):259-+ doi:10.1038/nchembio.1476

Sakr AA, Ghaly M, Ali M, Abdel-Haliem M (2013) Biodeterioration of binding media in tempera paintings by Streptomyces isolated from some ancient Egyptian paintings. Afr Jl of Biotechnol 12(14):1644

Santala S, Efimova E, Koskinen P, Karp MT, Santala V (2014) Rewiring the Wax Ester Production Pathway of Acinetobacter baylyi ADP1. Acs Synth Biol 3(3):145–151 doi:10.1021/sb4000788

Schlossman D, Shao Y (2014) Natural ester, wax or oil treated pigment, process for production thereof, and cosmetic made therewith. Google Patents

Simpson RJ (2007) Staining proteins in gels with coomassie blue. CSH protocols 2007:pdb. prot4719

Studier FW (2005) Protein production by auto-induction in high-density shaking cultures. Protein expression and purification 41(1):207–234

Trani M, Ergan F, Andre G (1991) Lipase-Catalyzed Production of Wax Esters. J Am Oil Chem Soc 68(1):20–22 doi:10.1007/Bf02660302

Tsujita T, Sumiyoshi M, Okuda H (1999) Wax ester-synthesizing activity of lipases. Lipids 34(11):1159–1166 doi:10.1007/s11745-999-0467-4

Tulloch A (1971) Beeswax: structure of the esters and their component hydroxy acids and diols. Chemistry and Physics of Lipids 6(3):235–265

USDA (U.S. Department of Agriculture) NASS (2017) Poultry Slaughter - 2016 Summary. In: USDA (ed). 24 February 2017 edn. USDA, p 1–35

Walker JM (2009) The Bicinchoninic Acid (BCA) Assay for Protein Quantitation. Springer Protoc Hand:11–15 doi:10.1007/978-1-59745-198-7_3

Wertz PW, Stover PM, Downing DT (1986) A Survey of Polar and Nonpolar Lipids from Epidermis and Epidermal Appendages of the Chicken (Gallus-Domesticus). Comp Biochem Phys B 84(2):203–206 doi:10.1016/0305-0491(86)90206-3

Yamada C, Sawano K, Iwase N, Matsuoka M, Arakawa T, Nishida S, Fushinobu S (2016) Isolation and characterization of a thermostable lipase from Bacillus thermoamylovorans NB501. J Gen Appl Microbiol 62(6):313–319 doi:10.2323/jgam.2016.06.002

Zhang Y, Meng K, Wang Y, Luo H, Yang P, Shi P, Wu N, Fan Y, Li J, Yao B (2008) A novel proteolysis-resistant lipase from keratinolytic Streptomyces fradiae var. k11. Enzyme Microb Tech 42:346–352 doi:10.1016/j.enzmictec.2007.10.015

